# Efficient genome editing of *Magnetospirillum magneticum* AMB-1 by CRISPR-Cas9 system for analyzing magnetotactic behavior

**DOI:** 10.1101/284786

**Authors:** Haitao Chen, Sheng-Da Zhang, Linjie Chen, Yao Cai, Wei-Jia Zhang, Tao Song, Long-Fei Wu

## Abstract

Magnetotactic bacteria are a diverse group of microorganisms with the ability to use geomagnetic fields for direction sensing. This magnetotactic behavior can help microorganisms move towards favorable habitats for optimal growth and reproduction. Highly efficient genomic editing is very useful for a comprehensive understanding of the magnetotactic mechanism at the molecular level. In this study, we adapted an engineered CRISPR-cas9 system for efficient inactivation of gene in a widely used magnetotactic bacteria model strain, *Magnetospirillum magneticum* AMB-1. By combining an engineered nuclease-deficient Cas9 and single-guide RNA, a CRISPR interference system was successfully developed to silence *amb0994* expression. More importantly, we succeeded in the construction of a single *amb0994* gene deletion mutant using CRISPR-Cas9 with approximate 60-fold high efficiency compared to classical homology double-crossing replacement procedure. This mutant synthesized normally the magnetosomes, but reacted quicker and with less time than the wild-type strain to abrupt magnetic field reversals. A dynamics simulation by modeling *M. magneticum* AMB-1 cell as an ellipsoid showed that the difference of the motions between wild and *Δamb0994* is due to flagellar influence. The behavior observation being consistent with dynamics simulation indicated that Amb0994 is involved in the cellular response to magnetic torque change via controlling flagella. Besides the contribution to a better understanding of the magnetotaxis mechanism, this study demonstrates the CRISPR system as a useful genetic toolbox for high-efficiency genome editing in magnetotactic bacteria.

## 1. Introduction

Magnetotactic bacteria (MTB) are a diverse group of prokaryotes that are able of sensing and changing their orientation in accordance with geomagnetic fields, a behavior known as “magnetotaxis”. This behavior is thought to facilitate the dwelling of bacteria within growth-favoring water columns or sediments with vertical chemical stratification [1, 2]. This unique capability is facilitated by special organelles that are synthesized intracellularly and membrane-enclosed ferromagnetic nanocrystals of magnetite (Fe_3_O_4_) and/or greigite (Fe_3_S_4_), called magnetosomes [3, 4]. Magnetosomes have emerged as a model for investigating prokaryotic organelle formation and biomineralization. However, the molecular mechanism of magnetotaxis in MTB is not understood in detail. The ability to precisely manipulate genomes is highly desirable in applications ranging from genetic analysis of functional genomic loci to the mechanisms of magnetotaxis. Current methods in the genetic modification of MTB models are classical homologous recombination (HR) and transposon mutagenesis [5-8]. Unfortunately, these approaches have low efficiency in precise sequence replacement [9].

Clustered regulatory interspaced short palindromic repeats (CRISPR) and CRISPR-associated (Cas) proteins based genome editing system has been developed [10, 11]. The type II CRISPR-Cas9 from *Streptococcus pyogenes* consists of only two elements, an endonuclease Cas9 and engineered single-guide RNA (sgRNA) [12]. Guided by a protospacer-adjacent motif (PAM) and a 20 nucleotide (nt) sequence matching the protospacer of the sgRNA, the Cas9-sgRNA ribonucleoprotein (RNP) complex binds specifically to the DNA target by sequence complementarity and induces a double-stranded breaks (DSBs) [12, 13]. A modified method, derived from the *S. pyogenes* CRISPR, CRISPR interference (CRISPRi) has been developed. Using an engineered nuclease-deficient Cas9, termed dCas9, that enables the repurposing of the system for the targeted silencing of transcription [14-17]. To date, CRISPR system has been successfully applied to a wide variety of eukaryotic [18-22]. To repurpose this system for genome engineering in prokaryotic cells, researchers use a homology repair donor to efficiently generate mutations with the assistance of the CRISPR-Cas9 DNA cleavage system, so that high-efficiency genomic deletions can be performed via homology-directed repair (HDR) integration [23]. Recently, many groups have worked on the adaptation of CRISPR-Cas9 system to bacteria, such as *Escherichia coli* [23], *Streptococcus pneumoniae* [23], *Lactobacillus reuteri* [24], *Streptomyces species* [25, 26], *Tatumella citrea* [27], *Clostridium genus* [28] and *Bacillus thuringiensis* [29], and so on. *M. magneticum* AMB-1 serves as a model organism in MTB for studying biomineralization and magnetotaxis. Although basic genetic tool as HR is available for MTB, precise gene modifications of HR rely on the integration of double-crossover events, the efficiency is low to enable full exploration and exploitation for its function [5, 6]. Compared with HR, it is likely that the performance of CRISPR is high recombinogenic capability in the host strains, and also offers unprecedented modularity [25, 26].

A large number of microorganisms respond to their physiological needs through the motion in their physicochemical microenvironments. The motion of bacteria triggers a change of swimming behavior via sensing extracellular stimuli, referred to as chemotaxis, phototaxis, and aerotaxis [30, 31]. In particular, magnetotaxis is found in MTB and is defined as the passive alignment of the cells to the geomagnetic field along with active swimming [32]. A model has suggested that magnetotaxis, together with aerotaxis, enables the MTB to move to suitable environmental conditions; this behavior is called “magneto-aerotaxis” [33]. However, several researchers think that magnetotactic behavior results from the active sensing of magnetic force to some degree [34, 35]. The active sensing process is a signal transduction system that depends on transmembrane chemoreceptors, known as methyl-accepting chemotaxis proteins (MCPs). These MCPs in turn could relay a signal to the flagellar motor switch protein, through a CheA-CheY signal transduction system [36, 37]. Interestingly, *M. magneticum* AMB-1 contains a large amount of MCP-coding genes [38]. As reported, Amb0994 is proposed as a special MCP-like protein in *M. magneticum* AMB-1 that interacts with MamK and plays a key role in magnetotaxis [37, 39]. Overproduction of Amb0994 results in slow cellular response to change along the direction of the magnetic field [37]. Moreover, Zhu et al. constructed a *Δamb0994-0995* double gene knockout mutant and studied its reaction to magnetic field changes. Results suggested that the cell senses the torque by Amb0994 and actively regulates the flagellar bias accordingly to align its orientation with the external magnetic field [40]. However, the construction of a single *amb0994* mutant has not succeeded in providing direct evidence to the involvement of Amb0994 in the active response to changes in the magnetic field. Therefore, we purposed as a proof-of-concept we studied the involvement of Amb0994 in magnetotaxis because the gene is involved only in magnetotactic behavior that is easy to observe.

In this study, we aim to explore a CRISPR-Cas9 system to develop a genomic editing platform for the magnetotactic model strain *M. magneticum* AMB-1. We successfully constructed *amb0994* knockdown (KD*amb0994*) and knockout strains (*Δamb0994*). Compared with current HR approaches, our CRISPR-Cas9 method has a higher editing efficiency. Analysis of the swimming behavior of the mutants showed the requirement of the Amb0994 MCP protein in the active response of the cells to magnetic field changes. Methods and principles we developed in this paper will provide useful reference for facilitating versatile and high-efficiency gene knockdown and deletion in MTB or other microorganisms.

## 2. Materials and Methods

### 2.1. Microorganisms and growth conditions

*E. coli* TOP10 was used for cloning and gene expression. WM3064 of *E. coli* was used as a donor strain in the conjugations and grown in the presence of 300 μM diaminopimelic acid (DAP). A concentration of 50 μg/mL of kanamycin (Kan) and gentamycin (Gm) were used in the *E. coli* culture. Cultures of wild-type and mutant strains of *M. magneticum* AMB-1 (American Type Culture Collection 700264) (Matsunaga et al. 2005; Matsunaga et al. 1991) were grown microaerobically in modified enriched *Magnetospirillum* growth medium (EMSGM) containing per liter: 5 mL of Wolfe’s mineral solution, 10 mL of Wolfe’s vitamin supplement, 5 mL of 0.01 M ferric quinate, 0.68 g of potassium phosphate, 0.12 g of sodium nitrate, 0.07 g of sodium acetate, 0.74 g of succinic acid, 0.05 g of L-cysteine–HCl, 0.2 g of polypeptone, and 0.1 g of yeast extract [41, 42]. The pH was adjusted to 6.75, 7 g of agar was added per liter of EMSGM to prepare the agar plates, and the medium was autoclaved. Most experiments were performed using a 100 mL Schott flask bottle or a 15 mL headspace (Providing 10% air in the headspace for 2% oxygen) and incubated at 30 °C. Generally, colonies appeared at the 6th–8th day after plating. Kanamycin was added at 15 μg/mL to agar plates and 10 μg/mL to liquid cultures. Gentamycin was added at 5 μg/mL.

### 2.2. DNA manipulation, plasmid construction and Conjugation

The enzymes for DNA modification were purchased from New England Biolabs (NEB) and Takara Biomedical Technology. Q5 High-Fidelity Polymerase (NEB) was used for PCR amplifications, except for colony PCRs, which were performed using PrimerSTAR HS DNA polymerase (Takara). All PCR products and plasmids were purified using Monarch DNA Gel Extraction Kit (NEB) and MiniBEST Plasmid Purification Kit (Takara), respectively. All plasmids and primers used in the study are listed in Tables S1 and S2.

#### CRISPRi-based *amb0994* suppression

Plasmid pCRISPRi-sgRNA*luxA* carrying dCas9, driven by the aTc-inducible TetR promoter, Kan^R^ was constructed as previous description [43]. In *M. magneticum* AMB-1, the TetR promoter was placed with the *tac* strong promoter under the control of an isopropyl-beta-D-thiogalactopyranoside (IPTG). The sgRNA is a 102-nt-long chimeric noncoding RNA, consisting of a 20-nt target-specific complementary region, a 42-nt dCas9-binding RNA structure, and a 40-nt transcription terminator derived from *S. pyogenes*. We designed *amb0994* as an sgRNA to the nontemplate (NT) DNA strand of the protein-coding region that blocks transcription elongation. By using software packages sgRNAcas9 and NCBI blast, we designed CRISPR sgRNA and potential off-target cleavage sites were evaluated [44]. The gRNA sequences were as follows: *amb0994*: 5′-GTAATATCGACCATGATTGG-3′. The sgRNA was changed using the primers sgRNA-0994-F and sgRNA-0994-R. The method was formulated by Larson [15]. We successfully constructed plasmid pCRISPRi-sgRNA*amb0994* and control plasmid pCRISPRi-no sgRNA. All constructions were checked by PCR, digestion, and sequencing.

#### Generation of *Δamb0994*

Two methods were used to create a deletion of *amb0994*. First, *Δamb0994* was constructed by HR. An approximately 1.0 kb upstream and downstream region flanking of *amb0994*, and the gentamycin resistance cassette from pUCGm were amplified using primers (Table S2). Amplified DNA fragments were ligated into the pMD18-T cloning vector (Takara) and cut with appropriate restriction enzymes; subsequently, they were cloned into the suicide vector pUX19 to form pUXsuc0994 [45]. Second, *Δamb0994* was constructed by CRISPR-Cas9 system-assisted HDR. Plasmid pCRISPR-sgRNA *amb0994* was constructed based on plasmid pCRISPRi-sgRNA*amb0994* from the following fragments: promoter tac; Cas9 was synthesized by a fusion PCR reaction that was performed with four primers (Primers Cas9-*XhoI*-F1, Overlap Cas9-R2, Overlap Cas9-F3 and Cas9-*XhoI*-R4) for gene mutation at two loci in dCas9; sgRNA (*amb0994*) was synthesized by GeneWiz inc.; similarly, approximately 2.0 kb HDR DNA fragment, including the 1 kb left and right arms of editing template, was amplified from purified genomic DNA of *M. magneticum* AMB-1, which inserted a gentamycin resistance cassette as a selectable marker. Correct plasmid assembly was confirmed by PCR, digestion, and sequencing (Sangon biotech).

#### Complementation of *Δamb0994*

Plasmid pAK0994 was assembled to complement *Δamb0994*, named com*amb0994*, a pBBR1MCS-based plasmid carrying a kanamycin resistance gene and expressing the *amb0994-*GFP fusion from a *tac* promoter. The method was performed as described previously [46]. *amb0994* was PCR amplified from *M. magneticum* AMB-1 with the primers Com*amb0994*-*EcoRI*-GFP-F and Com*amb0994*-*BamHI*-GFP-R. Amplified DNA fragments were digested by *EcoRI* and *BamHI* and ligated into plasmid pAK20 to create pAK0994. Relative abundance of Amb0994 in the cells was checked with a fluorescence microscope. Statistical results showed that approximately 40% of the cells did not express the protein (Data not shown). This phenomenon might be due to the stability of the plasmid or the amount of expression from the *tac* promoter [46].

Conjugation experiments both HR and CRISPR were performed as described previously with slight modification of the culture condition [7, 37]. WM3064 cells carrying each plasmid were growed at 37 °C in LB medium, until OD600 reached 0.5. 500 μL of WM3064 culture was washed twice in LB medium and suspend the cells with 200 μL LB medium with DAP. Mixed the WM3064 culture with 20ml of AMB-1 culture that were concentrated to a final volume of 200 μL and spot onto an EMSGM plate with DAP. The plate was incubated for 4 h at room temperature and the conjugations were suspended with 4 mL of EMSGM medium. The conjugations were then plated onto antibiotic-containing EMSGM plates and incubated in a microaerobic jar at 30 °C for 6–8 days. Following conjugation, single colonies were grown and checked for the presence of the interference construct. Resulting resistant strains were screened by PCR to determine the knockdown and knockout strains. This strain was checked for the absence of *amb0994*. The deletion region was sequenced to ensure that proper recombination occurred. In addition, six primers (p1, p2, p3, p4, p5, p6) were designed to verify the maintenance of magnetosome island (MAI) genes (Table S2). Six segments were chosen as *mamC, masA, mamB, mamY, mam O*′, and *mamK.* Clearance of the plasmid pCRISPR was performed on the plate without antibiotic to confirm restoration of kanamycin sensitivity.

### 2.3. Western blot analysis of *M. magneticum* AMB-1

*M. magneticum* AMB-1 colonies were transferred into 1.5-mL microcentrifuge tubes containing 1.5 mL EMSGM with 10 μg/mL kanamycin (For plasmid-bearing strains). A 1:10 dilution of *M. magneticum* AMB-1 cells in 1.5 mL of each culture was inoculated into 15 mL of EMSG media with kanamycin. Cultures were sampled at the same growth stage and were run in technical replicates. Cultures were grown in approximately 2% O_2_ for 1 day; after incubation for 24 h, 0.5 mM IPTG was added into half of the cultures to induce expression of the genes encoded by the plasmids and incubated at 30 °C for the next 24 h. Finally, cells were harvested by centrifugation at 10,000 rpm for 8 min at low temperature of 4 °C for Western blotting analysis.

The procedure for Western blotting was based on a general protocol as previously described [47]. Pelleted cells were resuspended in 20 mM Tris-HCl buffer (pH 7.4) and 5× protein-dyed buffer. All samples contained roughly equal amounts of cell by measuring cell density by spectrophotometer. Cells were heated at 100 °C for 10 min and loaded onto a 7.5% polyacrylamide gel, which was first run at 80 V for 0.5 h, and then at 100 V for 1 h to finish. Proteins were transferred onto a 0.22 μm polyvinylidene difluoride (PVDF) membrane at 100 V for 80 min. After the membrane was blocked for 1 h at room temperature with 5% milk-TBST (Tris-buffered saline-Tween 20), primary antibody (CRISPR-Cas9 monoclonal antibody recognizes both Cas9 and dCas9, Epigentek) was applied (1:500) for 1 h at room temperature. The PVDF membrane was rinsed thrice for 10 min with TBST, and secondary antibody (Goat anti-mouse conjugated with horseradish peroxidase [HRP], Thermo Fisher Scientific) (1:3,000) was applied for 1 h at room temperature. After washing with TBST thrice, antibody–protein complexes were then detected using Pierce CN/DAB substrate kit (Thermo Fisher Scientific, 34000), according to the manufacturer’s instructions.

### 2.4. Transmission electron microscopy (TEM)

For TEM observations, *M. magneticum* AMB-1 cells were first suspended in water, drop-cast, and dried onto carbon-coated copper grids. All TEM were performed on a JEM-2100 TEM with an accelerating voltage of 200 kV, equipped with an Oxford SDD detector (X-MaxN80T). For each experiment, at least 400 magnetosome crystals for 80 cells were imaged, and crystal size was measured by hand using the measuring tool in the software ImageJ 1.51i.

### 2.5. CRISPRi-based regulation of gene expression analysis

A total of 50 mL of each culture was centrifuged (8000×*g*, 10 min, 4 °C) and cell pellets were resuspended by vortexing in 1 mL of a mix of RNA protect solution and fresh culture medium (2:1). After 5 min incubation at room temperature, the cells were pelleted at maximum speed and flash frozen in liquid nitrogen for storage at −80 °C. Total RNA was extracted from cellular pellets with the RNAeasy Plus Minikit (Qiagen), which was quantified by spectrophotometry (NanoPhotometer). Contaminant DNA was removed from the RNA samples by digestion with gDNA Eraser. cDNA was produced from the RNA template by reverse transcription using the PrimeScript™RT reagent Kit with gDNA Eraser according to the manufacturer’s instructions (Takara), and stored at −80 °C for the next experiment. Reference genes are used to eliminate sample-to-sample variation. The RNA polymerase sigma factor *rpoD* was selected as a reference to normalize the data [47]. Primers are listed in Table S2, as designed by Primer 6. RT-qPCR was performed by following the manufacturer’s instructions for a SYBR green real-time PCR mix using an ABI StepOne Real-time Detection System (Applied Biosystems). The 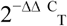 method was useful in the analysis of gene expression data [48]. All the samples from three independent experiments (Biological replicates) were analyzed in triplicate (Technical replicates), with negative controls included in each assay.

### 2.6. Magnetospectrophotometry assay analyses

Magnetospectrophotometry system was composed of a spectrophotometer (HITACHI U-2800, Japan) and an electromagnetic system [49]. Optical density (OD) and magnetic response (Cmag) of exponentially growing cultures were measured photometrically at 600 nm under the 4.5 mT homogeneous magnetic field, as described previously [50]. To avoid movement in the detection, 10 μM *m*-chlorophenylhydrazone (CCCP), a chemical inhibitor of oxidative phosphorylation, was added into the tube to stop cells from moving.

### 2.7. Motility analysis of *M. magneticum* AMB-1 by experiment

*M. magneticum* AMB-1 motility behavior was analyzed using a microscope equipped with custom-made electromagnetic coils [51]. Cells were subjected to a magnetic torque that reversed the direction of movement, resulting in an approximate U-trajectory after following field reversals, defined as “U turn” [52]. Swimming velocities, diameters, and times of “U turn” were determined using the MtrackJ plugin for ImageJ [53, 54]. Values of magnetic moment for cells were estimated from “U turn” analyses and from TEM of the magnetite chain by Esquivel and De Barros. The “U turn” method calculates the magnetic moment of a cell from its response time after a sudden magnetic field reversal. The cells were placed on a glass slide or in µ-Slide microchambers (Ibidi). To observe the glass slides, 4 µL of cells was observed under upright Olympus microscope using a long working distance ×40 objective and recorded at 33 fps using a sCMOS camera (AndorNeo). Otherwise, 75 μL of cell suspension was placed in a microchamber and observed with a ×40 phase-contrast microscope (Olympus), recorded at 20 fps using a Canon camera. In addition, the magnetic field was 1 mT in the “U turn” experiment without shielding the geomagnetic field. A total of ∼300 traces of ∼80 bacteria were analyzed in the“U turn” experiment, but ∼30 bacteria were analyzed in alpha angle trajectory.

### 2.8. Motility analysis of *M. magneticum* AMB-1 by simulation

We simulated the motion trajectory of bacteria in a magnetic field using an ellipsoidal model. The simulation was based on the concept that the moving *M. magneticum* AMB-1 cells, which are similar to ellipsoid, would suffer resistance in the fluid. The orientation of MTB under a magnetic field could be described by a partial differential equation:

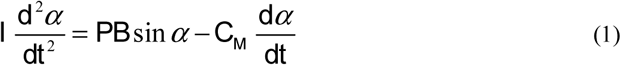

where *P* is the magnetic moment of one *M. magneticum* AMB-1, *B* is the applied magnetic field, and α is the angle between the direction of the magnetic field and the magnetic dipole moment. For *M. magneticum* AMB-1, the direction of magnetic dipole moment was the same as the major axis of the cell body, thus α could be measured by the angle between the instantaneous velocity vector and magnetic field line.

Other variables were defined as the following:

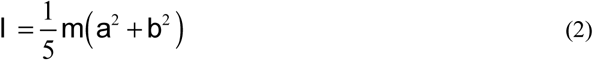

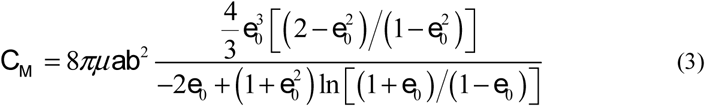

where *I* is the product of inertia of *M. magneticum* AMB-1, *m* is the mass, and *a* and *b* are the halves of the length and width of a *M. magneticum* AMB-1 cell, respectively. 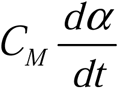 represents the resistance torque applied to the *M. magneticum* AMB-1 cell. *C*_*M*_ is the coefficient of the resistance torque. In the expression (3) of *C*_*M*_, *μ* is the viscosity coefficient of the culture medium, and *e*_0_ is the auxiliary variable, which is defined as 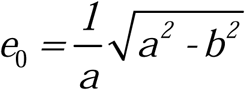.

Using the derivation method by Esquivel and De Barros, we can obtain the following expression of the alpha angle [52]:

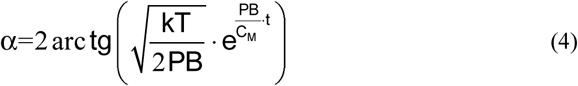

where *k* is the Boltzmann’s constant (1.38×10^−23^), *T* is the absolute temperature (K), and *t* is time.

Using the expression of (4), we calculated the orientation of *M. magneticum* AMB-1 under a magnetic field. If the bacterium has no flagella, or if the influence of flagellar movement and resistance in the fluid is not considered, the *C*_*M*_, defined as the expression of (3), could be utilized. If the bacterium has a flagellum, and flagellar influence suggesting a flagella movement signal provided by cells is considered, a modified *C*_*M*_ should be used. According to the research of Steinberger et al., when the *M. magneticum* AMB-1 is modelled as an ellipsoid connected with a wire fixed to one end of its body, the *C*_*M*_ should be multiplied by 4.67 (± 0.47) [55]. In our work, *C*_*M*_ was multiplied by 5 based on the expression of (3) for the case to consider the influence of flagellar movement and resistance in the fluid.

### 2.9. Statistical analysis

All statistical analyses were performed using SPSS 22.0 (SPSS, IBM). Two-tailed Student’s *t*-test was used in transcription level analysis. One-way analysis of variance was used to investigate the difference in velocities with magnetic field increase. Size, shape factor, and number of magnetosomes in one cell were analyzed using Mann–Whitney *U*-test. Each experiment was repeated thrice, and all data are expressed as mean ± SD. Level of significance of the differences observed between the control and test samples were expressed as one, two, or three stars, for \**P*<0.05 and \*\**P*<0.01. A *P* value of <0.05 was considered significant in all statistical tests.

## 3. Results

### 3.1 CRISPRi-mediated *amb0994* gene knockdown in *M. magneticum* AMB-1

CRISPRi system can be used to knockdown gene expression in eukaryotes and prokaryotes. To analyze the utility of CRISPRi for gene repression in *M. magneticum* AMB-1 cells, we first constructed an efficient dCas9 expression system for this strain. We chose to target NT strand of *amb0994*, as some researchers found effective transcriptional repression via NT-targeting of the gene [16]. The constructed plasmids were introduced into *M. magneticum* AMB-1 by conjugation as described in the “Materials and Methods” section. Western blot was performed to check the specific dCas9 expression. As shown in Figure 1a, anti-dCas antibodies specifically identified the 160 kDa polypeptide with the anticipated molecular weight for dCas9 protein in *E. coli* TOP10 and *M. magneticum* AMB-1 cells with CRISPRi plasmid, as well as the control strains (Plasmid without sgRNA). Further RT-PCR analysis showed that the dCas9 expression was significantly increased with IPTG induction in these two strains (Fig. 1b). Thus, IPTG was used in all experiments.

**Fig. 1.**
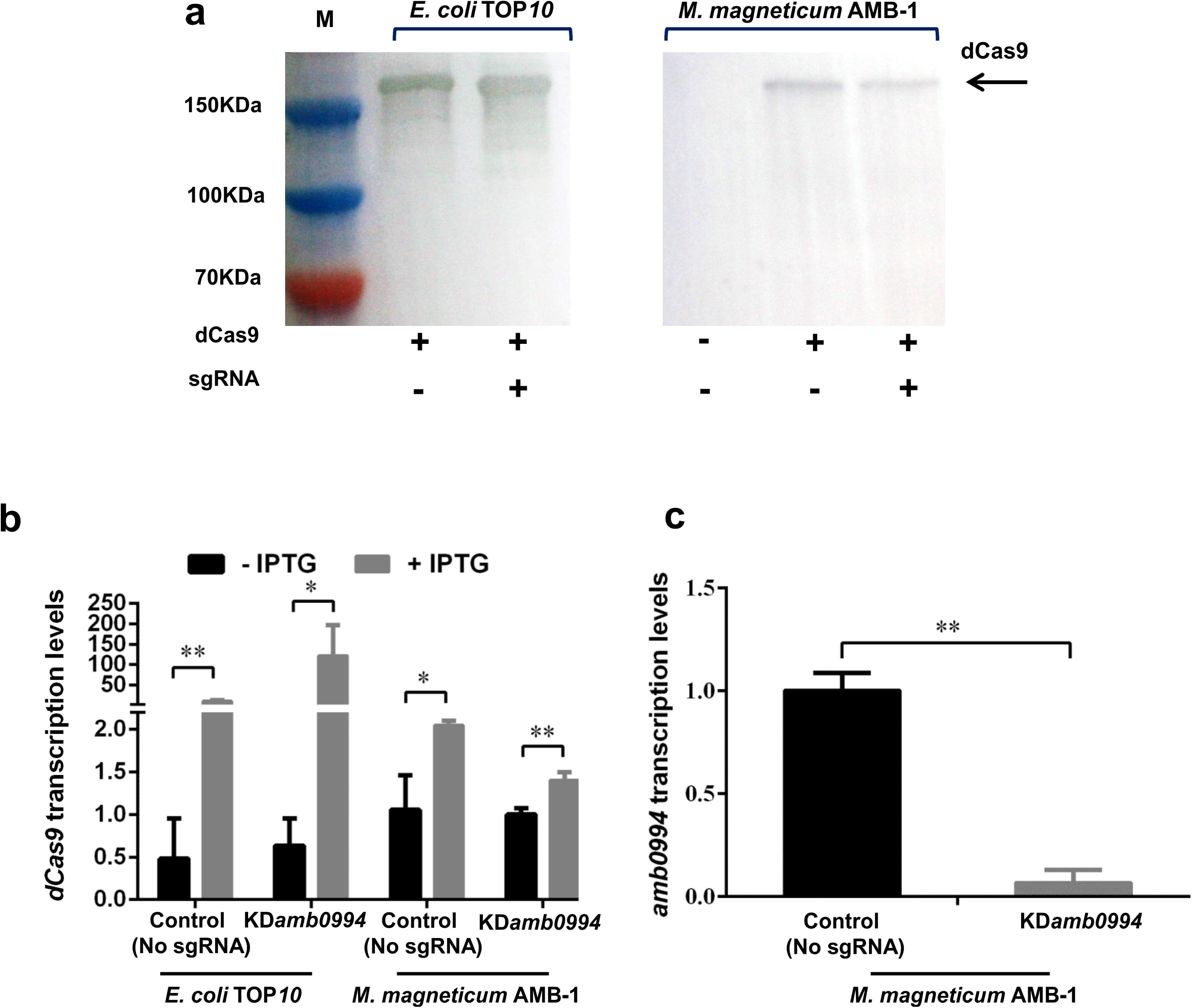
CRISPRi efficiently represses the *amb0994* gene transcription. (a) Western blot detection of dCas9 after SDS-PAGE with whole cell samples from experiment. The M represents the positions on the gel of the molecular mass markers (from top to bottom: 150, 100 and 70 kDa). (b) *dCas9* transcription levels of the control (plasmid without sgRNA) and test strains in *E. coli* and *M. magneticum* AMB-1 cells (N=3 per strain) were determined. The error bars represent the standard deviation of samples. (c) The relative expression levels of *amb0994* in the KD*amb0994* compared to control strains without sgRNA. The levels of transcription were calculated relative to the housekeeping gene *rpoD* (N=9 per strain). The error bars represent the standard deviation of samples. \**P*<0.05, \*\**P*<0.01.

We analyzed the consequence of dCas9 production on the levels of transcription of the *amb0994* gene (Fig. 1c). The expression of a plasmid-borne dCas9 induced by IPTG in cells expressing gRNAs targeted at the NT strand of the *amb0994* gene resulted in *amb0994* repression. The relative levels of mRNA of *amb0994* were strongly reduced by nearly 93% (*P*<0.001) versus the control strain, which expressed dCas9 but lacked sgRNA. Therefore, the CRISPRi-dCas9 system was used to efficiently knockdown the *amb0994* expression.

### 3.2. Effient genome editing by CRISPR system compared with traditional HR

CRISPRi-dCas9 is an efficient approach for studying gene functions and for engineering genetic regulatory systems from sequence-specific control of gene expression on a genome-wide scale. However, off-target effects caused by CRISPRi technology might perturb gene expression at off-target sites. To further verify the function of Amb0994, we construct neat amb0994 deletion mutant. Previously reported CRISPR genetic modification technologie has been successfully applied to a wide variety of organisms. To assess the ability of Cas9-gRNAs in promoting homology-directed repair in *M. magneticum AMB-1 strains*, we further compared the efficiency of Cas9-gRNAs versus traditional HR in targeting mutation loci by measuring HDR donor-based precise gene editing.

We performed CRISPR-Cas9-mediated knockout experiment. By designed a HDR donor template with homology arms near the Cas9-gRNA, sgRNA transcripts could guide Cas9 nuclease to introduce DSBs at the ends of the *amb0994* gene, while a codelivered editing template repairs the gap via HDR (Figs. 2a and 2b). The resulting resistant strains were screened and evaluated by PCR from genomic DNA with primers that anneal outside of the 1 kb homology arms and the maker region of gentamycin resistance cassette. A total of 9 out of 15 (60%) randomly selected colonies confirmed the deletion of *amb0994*. As shown in Figure 2c, a PCR amplification with primers annealing with upstream of 1 kb lift arm (Primer 1) and gentamycin resistance cassette (Primer 3) was performed; An approximately 2.0 kb band indicative of deletion was observed in five representative colonies, whereas no band was observed when wild-type genomic DNA was used as the template. The result showed that the gentamycin resistance cassette has replaced the fragment that needs to be deleted. Similarly, the sequences of the deleted region were amplified by PCR with primers 1 and 2. The result showed a 1.0 kb band from the wild-type genomic DNA, but not from genomic DNA of the strains in which the *amb0994* were deleted (Fig. 2d). The PCR fragment was sequenced with internal primers to determine if the intended deletion was introduced. In the meantime, we designed a suicide plasmid of pUXsuc0994 for deletion through traditional HR as described in the “Materials and Methods” section. A total of 1 out of 101 randomly selected colonies confirmed the deletion of *amb0994*. The results showed that only 1% selected colonies were correctly edited for homologous double-crossover (Data not shown). We successfully use CRISPR-Cas9 system and HR methods to knockout the gene *amb0994* in *M. magneticum* AMB-1. Importantly, CRISPR system is approximately 60-fold more efficient than HR in construction of *amb0994* deletion mutants.

**Fig. 2.**
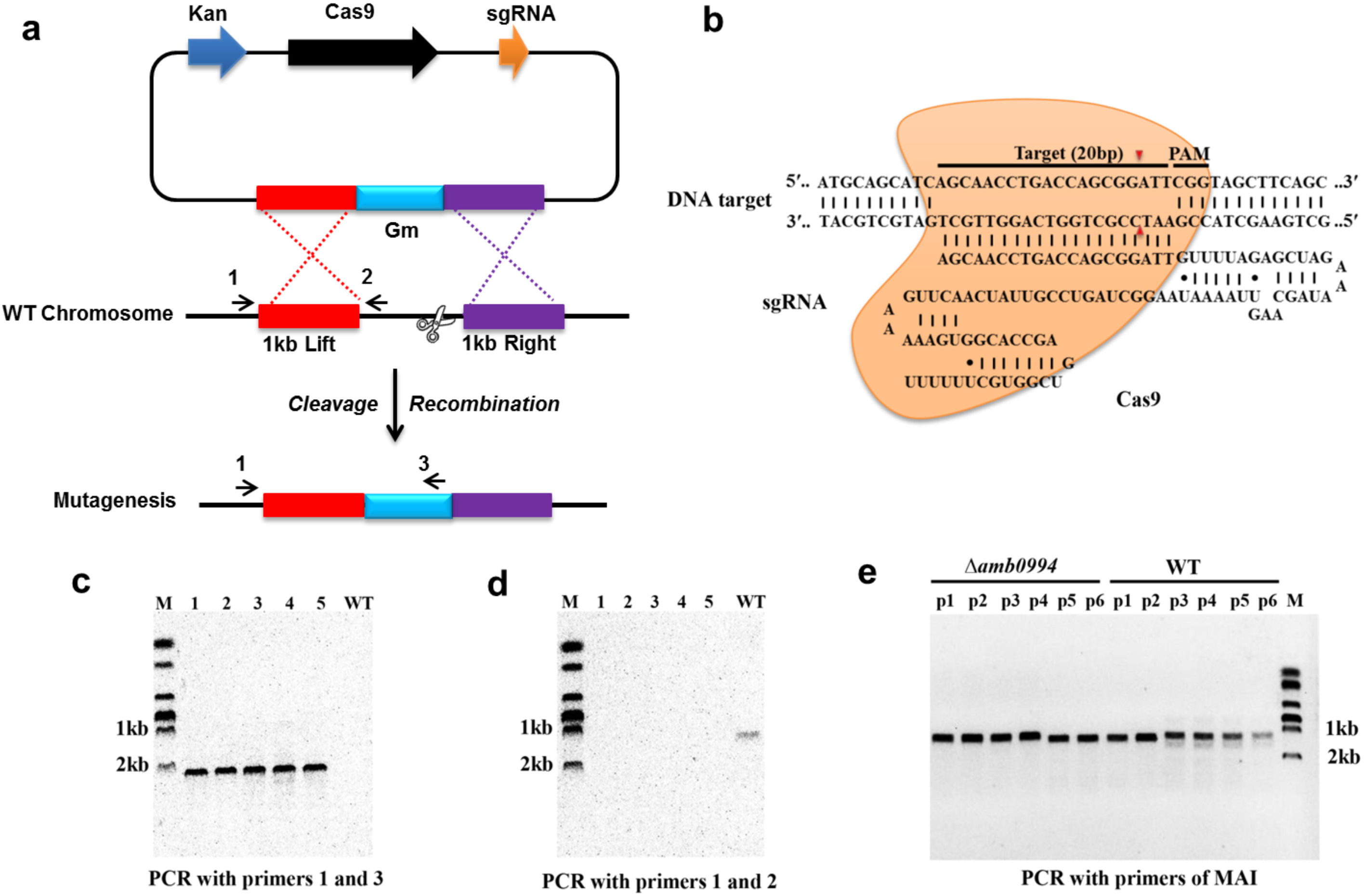
CRISPR-Cas9-assisted genome editing in AMB-1 cells. (a) Strategy for deletion of the *amb0994* gene by CRISPR-Cas9 assisted HDR in *M. magneticum* AMB-1 cells. An sgRNA transcripts guide Cas9 nuclease to introduce DSBs at ends of *amb0994* gene, while a codelivered editing template repairs the gap via HR. Kan is kanamycin. Gm is gentamycin. (b) Schematic of RNA-guided Cas9 nuclease uses for editing of the AMB-1 *amb0994*. An sgRNA consisting of 20 nt sequence (black bar) guide the Cas9 nuclease (orange) to target and cleavage the genomic DNA. Cleavage sites are indicated by red arrows for ∼3 bp upstream of PAM. (c and d) PCR evaluation of *amb0994* deletion from five colonies (1−5) with WT control. (e) Six fragments within MAI were amplified to evaluate the maintenance of genomic MAI during deletion.

In addition, MTB share a conserved genomic island, termed MAI, which is involved partly in magnetosome formation and encodes most of the proteins that are physically associated with purified magnetosomes [38]. MAI is genetically unstable and results in frequent spontaneous loss of the magnetic phenotype [56]. We designed six pairs of primers within MAI to evaluate the maintenance of genomic MAI during deletion. Result showed that the genomic MAI in the deletion strains of *amb0994* was not lost (Fig. 2e). Taken together, all these results showed that the advantage of using CRISPR-Cas9 with an HDR donor would afford higher editing efficiency by nearly 60% compared with using traditional homologous double-crossover (1%) in construction of *amb0994* deletion mutants.

### 3.3. Effect of Amb0994 suppression or deletion on magnetosome formation in *M. magneticum* AMB-1 strains

After constructing KD*amb0994* and *Δamb0994*, we checked the magnetosome synthesis by analyzing the Cmag of cells and the size, shape factor and number of magnetosomes in KD*amb0994, Δamb0994*, control, and Com*amb0994*. The *cas9* gene suggested an inherent toxicity associated with the overexpression of the heterologous endonuclease; thus, we chose the strain that transformed a plasmid without sgRNA but expressed Cas9 as a control to KD*amb0994*, whereas the wild-type *M. magneticum* AMB-1 (WT) without plasmid was used as the control to *Δamb0994* and Com*amb0994*.

In Figure 3a, *amb0994* suppression or deletion did not affect the value of Cmag versus the control. We analyzed the size, shape factor, and number of magnetosomes in *M. magneticum* AMB-1 cells according to TEM observations. Result showed that the sizes of the magnetosome were 40.08 ± 11.45 nm and 39.91 ± 12.74 nm for the control (No sgRNA) and KD*amb0994*, respectively (Figs. 3b_1 and 3c_1). The shape factors were 0.87 ± 0.09 and 0.89 ± 0.09 (Figs. 3b_2 and 3c_2). The numbers of magnetosomes in each cell were 21.33 ± 5.87 and 21.28 ± 4.18 (Figs. 3b_3 and 3c_3). Statistical analysis of U-test revealed insignificant differences between *amb0994* suppression cells and control (No sgRNA) (*P*>0.05). Moreover, we further analyzed the TEM data in *Δamb0994*, Com*amb0994*, and WT control. Indeed, the results of the size, shape factor, and number of magnetosomes in *M. magneticum* AMB-1 cells within these strains were the same as in the CRISPRi strains (Figs. 3d_1–3, 3e_1–3, 3f_1–3, and 3g_1–3). These results indicated that the suppression or deletion of *amb0994* in *M. magneticum* AMB-1 would not interfere with the synthesis of magnetosomes.

**Fig. 3.**
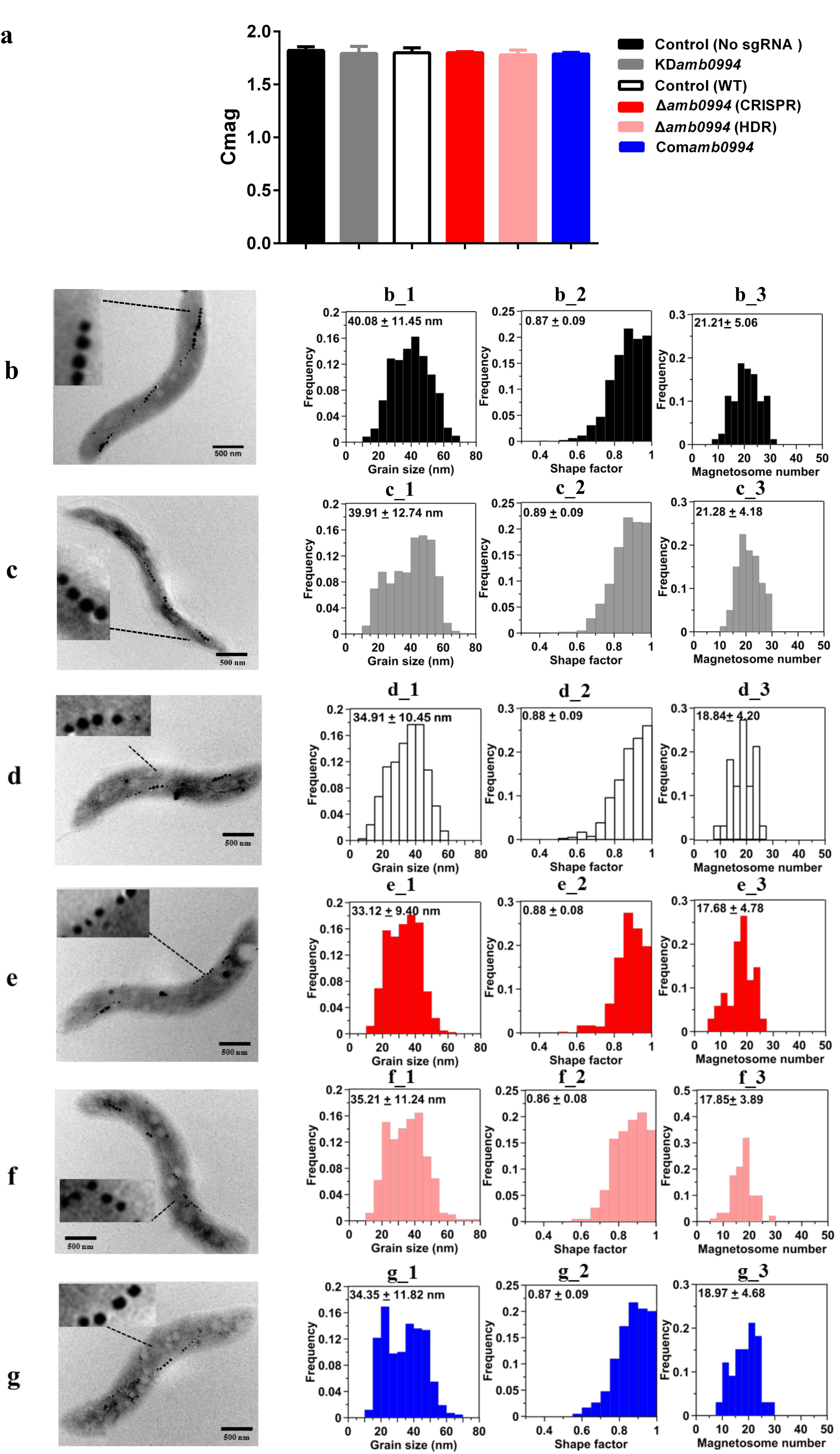
Magnetosome mineralization is not affected by CRISPR-mediated *amb0994* suppression or deletion in *M. magneticum* AMB-1. (a) Cmag curves of control, KD*amb0994, Δamb0994* and com*amb0994* strains under the 4.5 mT magnetic field, recording of AMB-1 treated with CCCP. TEM images of control strain without sgRNA (b), KD*amb0994* (c), WT control (d), *Δamb0994* by CRISPR (e), *Δamb0994* by HDR (f) and com*amb0994* (g). (b_1–3)–(g_1–3) Histogram analyses of the number, size, and shape factors of magnetosomes in these strains. The scale bar is 500 nm.

### 3.4. Amb0994 is involved in the response to the magnetic field

To verify the role of Amb0994, we analyzed the motility behavior of *M. magneticum* AMB-1 in a laboratory magnetic field by an optical microscope. Studies have shown that the motion trajectory of MTB is normally a “U turn” when a reversal magnetic field is applied. The swimming velocities, diameters, and times of “U turn” following field reversals were measured as described in the “Materials and Methods” section. After 48 h of culture, the cells were placed in microchambers or glasses with a uniform magnetic field applied.

Magnetic field strengths (0, 0.5, 1, and 1.5 mT) had no apparent effects on the velocities of KD*amb0994* cells and control strains, whereas the average swimming velocities of KD*amb0994* were significantly higher than those of the control (*P*<0.05) (Fig. 4a). More importantly, the diameters of “U turn” in the cells of KD*amb0994* under 1 mT magnetic field were smaller, and the times of “U turn” in KD*amb0994* were shorter compared with those of the control (Figs. 4b and 4c, *P*<0.05, Movies S1 and S2). These results indicated that the knockdown mutant reacted faster and spent less time to achieve the “U turn” in response to magnetic field reversal.

**Fig. 4.**
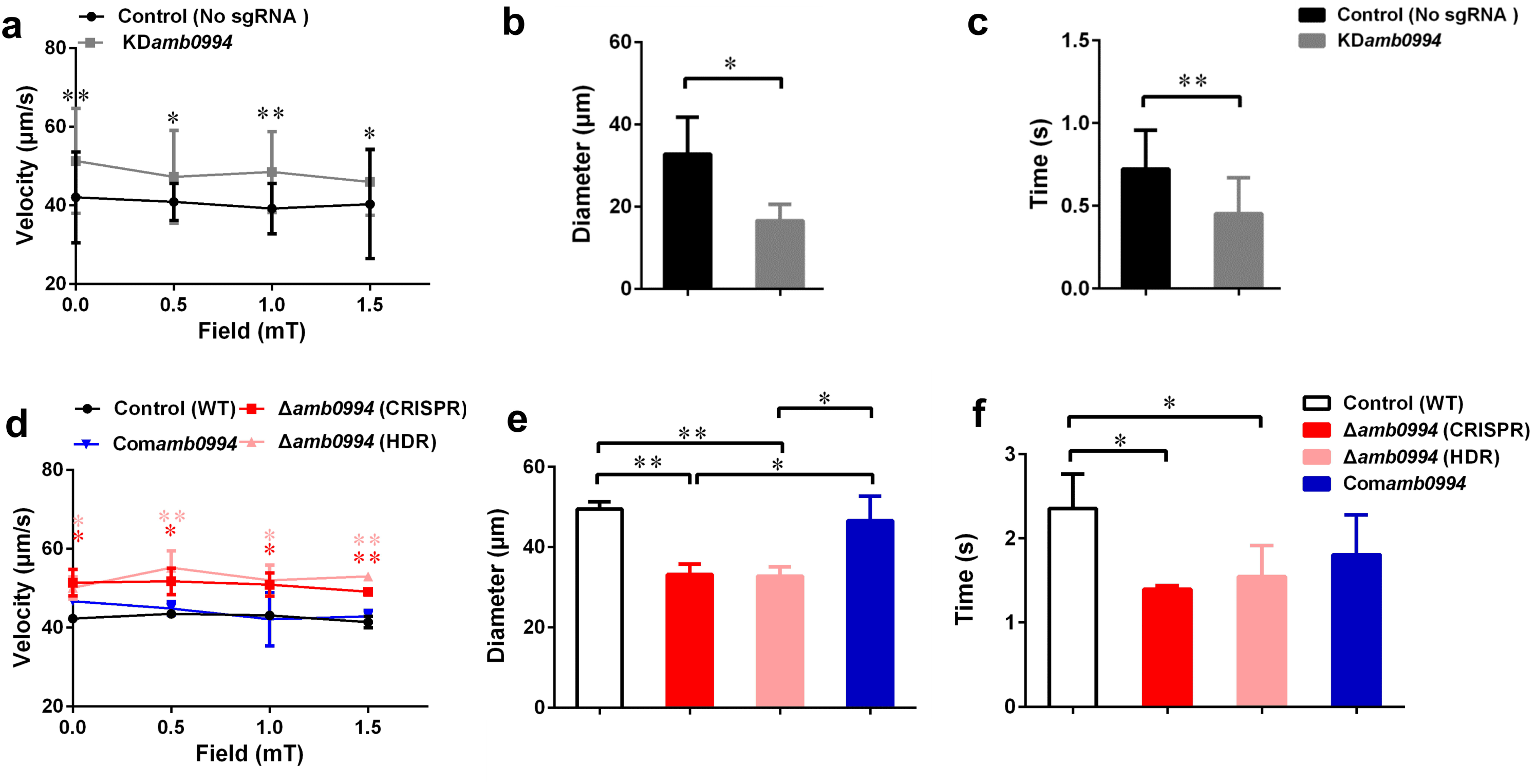
Effects of *amb0994* suppression or deletion on the swimming behavior of *M. magneticum* AMB-1 cells. (a and d) Swimming velocities were measured under different fields. Red stars represent the statistical analyses performed between *Δamb0994* (CRISPR) and WT control, whereas pink stars show statistical analyses between *Δamb0994* (HDR) and WT control. (b, c, e, and f) Diameters and times of “U turn” were analyzed in a 1 mT uniform magnetic field. Gray lines and bars represent the KD*amb0994* cells, black lines indicate control without sgRNA or wild-type groups, red and pink indicate *Δamb0994*, and blue bars show Com*amb0994* strains. \**P*<0.05, \*\**P*<0.01.

To corroborate the involvement of Amb0994 in response to magnetic field changes, we analyzed motility behavior of the neat *Δamb0994* deletion mutant, the corresponding complementation of Com*amb0994*, and WT strains. Average swimming velocities of CRISPR-mediated *Δamb0994* were significantly faster than those of the complemented strain and wild-type control (*P*<0.05) (Fig. 4d). In addition, compared with those of the control, the diameters and times of “U turn” in the cells of CRISPR-mediated *Δamb0994* were smaller and shorter, respectively (Figs. 4e and 4f, *P*<0.05). These results showed that the “U turn” of CRISPR-mediated knockout mutant was faster than that of wild-type cells. The same results were obtained with traditional deletion mutant. The behavior observation, together with CRISPR-mediated knockdown or knockout mutants, consistently implied that Amb0994 is involved in cellular responses to magnetic torque changes via controlling flagella.

### 3.5. Amb0994 suppression or deletion reveals a different alpha angle trajectory in “U turn”

Previous work found that active sensing exists in magnetotaxis and Amb0994 functions as a magnetic receptor that senses the instantaneous alpha angle (*α*) between the instantaneous velocity vector and the magnetic field line, which was used to reflect the angle between the cell body and magnetic field direction [40]. To research the magnetotactic behavior mechanism, we analyzed the alpha angle trajectory in “U turn” in KD*amb0994* and control (no sgRNA). Our experiments showed that the KD*amb0994* slop (gray line) was significantly sharper and the time was significantly shorter compared with that (black line) of the control (*P*<0.05) (Fig. 5). This result was consistent with “U turn” data and showed a smaller diameter and less time spent by knockdown *amb0994* strain.

**Fig. 5.**
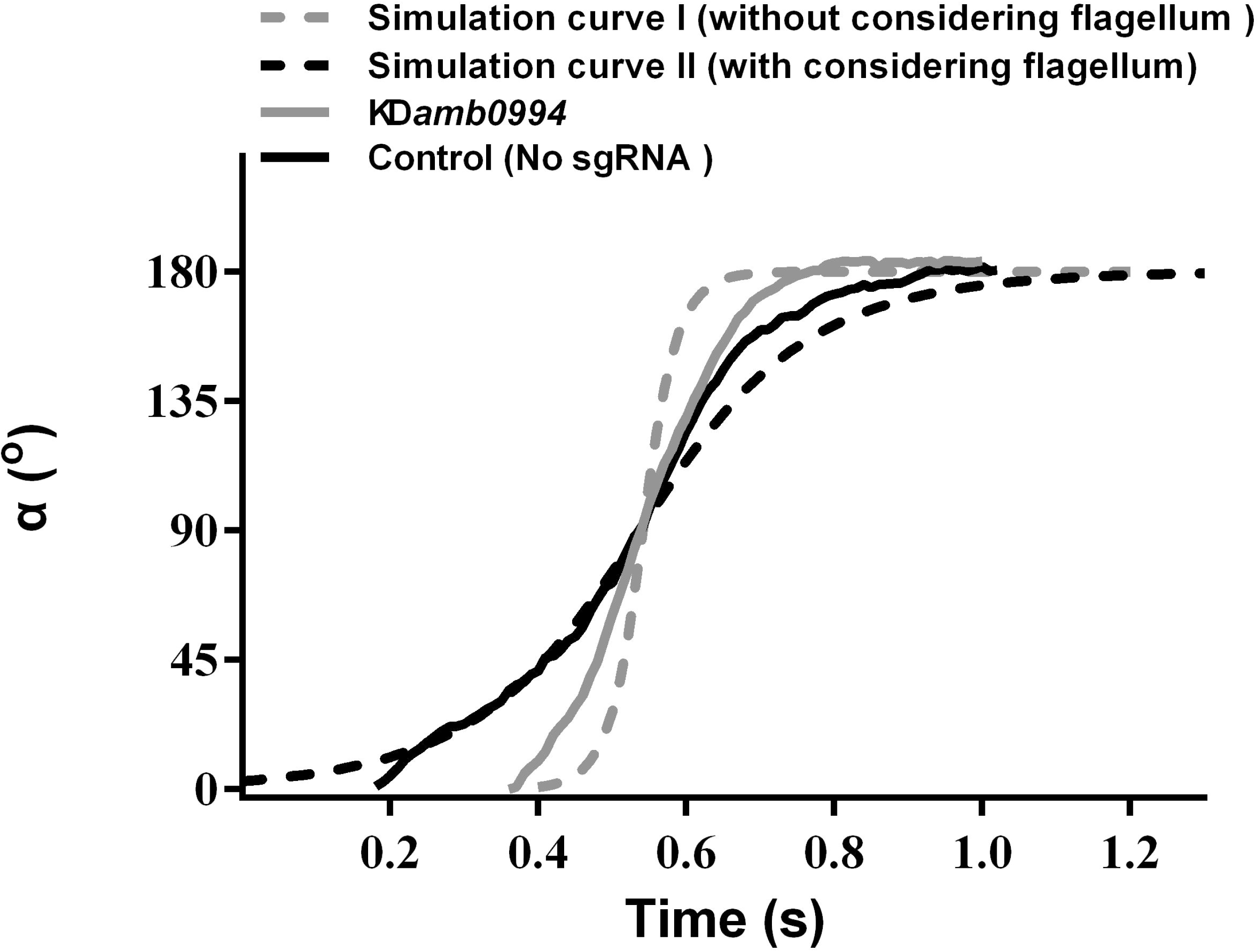
Trajectory of alpha angle after the magnetic field is reversed. The black solid line is the trajectory for the control cell. The gray solid line is for the CRISPRi-mediated silencing mutant of *amb0994* as KD*amb0994.* The black dash-line is the simulation curve with considering the effect of the flagellum. The gray dash-line is the simulation curve without considering the flagellum. To compare these curves, we assumed that 90° of the alpha angle occurred simultaneously. The curve is generated from the typical representative data of ∼30 bacterial cells.

To further explain this behavior, we simulated the motion trajectory of bacteria in a magnetic field using an ellipsoidal model with or without considering the flagellar effect. According to the sizes of *M. magneticum* AMB-1 cells, the ellipsoid was assumed as 3.2 μm (major axis diameter) ×0.5 μm (minor axis diameter) ×0.5 μm (minor axis diameter). For modeling *M. magneticum* AMB-1 cells without considering the effect of the flagella, we calculated the resistance factor *C*_*M*_ according to expression of (3) as described in the “Materials and Methods” section. To model *M. magneticum* AMB-1 cells with consideration of the flagella in the mathematical simulation, we chose the experimentally determined resistance factor as 5*C*_*M*_. As shown in Figure 5, the slope without considering the flagella (gray dash-line) is sharper, whereas the time is shorter compared with that (black dash-line) when the flagella is considered. The experimental curve of the KD*amb0994* was closer to the simulated curve without considering the flagellar effect, whereas the experimental curve of the control was closer to the simulated curve when considering the flagellar effect.

To validate the function of Amb0994, we analyzed the trajectory for *Δamb0994* and Com*amb0994*. Results showed that the motion curve of *Δamb0994* was closer to the simulation curve without considering the flagella (data not shown). All the results would suggest that there is a signal, which is transferred to the motors to adjust the flagella and the swimming pattern of the cell when a reversal magnetic field is applied, that indirectly links *amb0994* gene and flagella movement. We further confirmed Amb0994 is essential for magnetic sensing.

## 4. Discussion and conclusion

Genetic engineering of biological systems possesses significant potential for applications in basic science, medicine, and biotechnology [12]. Recent work on CRISPR-Cas system has renewed genetic modification [12]. As an efficient genome editing tool, it can be markedly easy to design and highly specific for gene editing in diverse organisms. Moreover, CRISPRi-dCas9 system has been broadly used in targeting and silencing specific genes without altering the DNA sequence, and it can potentially be adapted for transcriptional regulation on a genome scale in various organisms [14]. Prior to this work, although current tools for genome editing are available for MTB models, the efficient of these tools is still too low to enable full exploration and exploitation of this special group of organisms. Hence, this is useful to apply the CRISPR-Cas9 system in MTB to improve the efficiency of genome editing.

As all we known, a well-conserved MAI of MTB is essential for magnetosome formation, thereby causing the bacterium to passively migrate along the field as it swims [38]. Thus, most of the current studies are focused on characterizing the deletion mutants of the MAI genes to analyze the molecular factors and processes in magnetosome biogenesis. The functional analysis of genes that maybe related with an active magnetotactic behavior but do not interfere with magnetosome formation is relatively limited. *amb0994* is the gene that is likely to be involved in magnetotactic behaviors, but there is still no direct evidence and even a lack of single *amb0994* mutant to confirm its function. In this study, we first used the CRISPRi technology to specifically repress *amb0994* transcription in attempts to understand its function. We successfully co-expressed dCas9 and sgRNA in *M. magneticum* AMB-1 cells and suppressed *amb0994* expression up to 93% (Fig. 1). As the results showed that the dCas9 expression was significantly increased with IPTG induction in these two strains, but dCas9 expression in *M. magneticum* AMB-1 was significantly lower than those in *E. coli.* It could because: 1. Plasmid copy number is different in two strains; 2. different metabolism activity in two host strains and 3. *M. magneticum* AMB-1 might be more sensitive to dCas9 than *E. coli*. Although CRISPRi can achieve stringent suppression with several sgRNA targeting the same gene, this result might cause off-target effects produced by CRISPRi technology. Thus, to clearly understand the function of *amb0994* gene, we generated a deletion of *amb0994* and its complementation strain to analyze its phenotype in detail. By rationally designing sgRNA, we increased the rate of genome editing and shortened the time for subsequent screening of the correct mutants in *M. magneticum* AMB-1 cells when using CRISPR-Cas9 system compared with classic HDR. However, some of the ex-conjugants were mixtures of edited and wild-type cells in this population. Nevertheless, the observed efficient genome editing occurred. This finding provides a tool for fast and high-efficiency silencing or deletion of genes in the MTB model strain *M. magneticum* AMB-1. To complicate large-scale DNA fragment suppression or deletion, this method will be directed to employ multiple sgRNA targeting different regions to achieve tighter knockdown or knockout of genes to generate editing with high efficiency in MTB [57].

The TEM analyses for size, shape factor, number of magnetosomes in one cell, and Cmag measurements were similar to those carrying the sgRNA vector and those that do not. By adding CCCP to interrupt flagellar rotation, we can eliminate the influence of cell motion on Cmag detection. These findings indicated that *amb0994* suppression or deletion in *M. magneticum* AMB-1 do not affect the passive alignment of cells along the magnetic field lines and interfere with the synthesis of magnetosomes. In addition, these results further exclude the concern of potential off-target activity and toxic effect of Cas9 using CRISPR-Cas9 gene editing system. However, in different batches of cultivation strains, factors such as culture conditions and activity of cells might affect the magnetosome formation. Thus, we should choose the same time to cultivate strains with the control and experimental groups to analyze the data in each repeat experiment.

Furthermore, by analyzing the swimming behaviors of *M. magneticum* AMB-1, the swimming velocities of *amb0994* suppression or deletion strains were significantly faster than those of the control. This finding might imply a default basic velocity without regulation of the flagellar motor rotation in the absence of Amb0994 function. In other words, Amb0994 might fine-tune the flagellar rotation speed to adapt to environmental or magnetic disturbances. According to the analysis by Esquivel and De Barros, the diameters of “U turn” in the MTB cells under a reversed magnetic field should be proportional to their swimming velocities, whereas the times of “U turn” should not be related with their swimming velocities [52]. However, the experimental results showed that *amb0994* mutants reacted quicker compared with control, and the diameters of “U turn” of *amb0994* mutants were smaller than those of the control. By analyzing the dynamics of the *M. magneticum* AMB-1 model, we suggest that the reason for the smaller diameters and faster times of “U turn” of *amb0994* mutants in response to the reversed magnetic field are related to the influence of the flagella. For the wild-type *M. magneticum* AMB-1, the perfect connection between cell body and flagellum causes the cells to have higher resistance in the fluid, which may help withstand perturbations in the magnetic field. For *amb0994* mutants, the absence of Amb0994 function may affect the junction between the flagella and cell body, or decrease the capability of the flagella following the change of the cell body. According to the early model, the magnetosome chains of cells generate magnetic torque when they are not aligned well along the magnetic field lines. The MCP Amb0994 senses the magnetic torque by interacting with MamK, and transfers the signal to the flagellar motors [37]. However, the motion coordination between the cell body and flagella to respond to magnetic field changes is unclear. Our results suggest that Amb0994 is involved in cellular response to magnetic torque changes via controlling flagella.

Research has shown that cells keep active motility under low magnetic fields, so they would take more time to respond to changes in magnetic field [37, 58]. This phenomenon further explains why the cells possessing *amb0994* can be active in sensing the external fields to coordinate their motions, rather than being completely passive to orientation. Another study revealed that *M. magneticum* AMB-1 cells can sense magnetic field gradients and respond to them by reversing direction [59]. The author speculates that this difference in the field could be torqueing the cells and relaying this signal to Amb0994. Consequently, combining with theoretical and experimental analyses, we can further confirm the hypothesis that the MCP like-protein Amb0994, as an active magnetic sensor, transmits magnetic signals to the flagellar system to control the movement of the flagella. Bacteria use a widely studied chemotactic signaling pathway to sense various chemicals and physical stimuli, and orient their movements to make them a favorable environment. MCPs are a family of bacterial receptors that detect stimuli and affect the direction of flagella rotation through a signal transduction system [60]. These may be thought to help bacteria quickly seek favorable habitats in weak geomagnetic fields and this may help achieve better insights on the magnetic response mechanism.

In conclusion, we demonstrated that the type II CRISPR-Cas system of *S. pyogenes* can be reconstituted in *M. magneticum* AMB-1 cells to efficiently and precisely silence or delete genes involved in magnetotaxis. This study provides an efficient and specific genome targeting and editing platform, and might pave a new avenue for genetically engineering the magnetotactic model strain. Together the behavior observation and dynamics simulation showed that Amb0994 is involved in cellular response to magnetic torque change via controlling flagella. These imply that the active sensing of magnetic field plays a key role in magnetotaxis.

## Acknowledgements

This work was supported by the National Youth Foundation of China (Grant NO. 31300689), the National Key Research and Development Program of China (Grant NO. 2017YFC0108501), and the Research Project Funded by the Institute of Electrical Engineering, Chinese Academy of Sciences (Grant NO. Y650141CSA). We greatly appreciate Xu Tang for TEM observation and Junwei Du for magnetic field measurement. We thank Xiaoxiao Zhu, Pingping Wang and Changyou Chen with the help of useful discussion and suggestions. We are grateful to Ying Li and Xu Wang for the plasmids of pUX19 and pUCGm. We also thank the anonymous reviewers and the editor for their valuable comments.

## Conflict of interest

The authors declare that they have no competing interest.

