## Supplementary Materials for "Efficient genome editing of *Magnetospirillum magneticum* AMB-1 by CRISPR-Cas9 system for analyzing magnetotactic behavior"

**Table S1** Plasmids used in this study

| Plasmids | Relevant genotype | Reference |
|---|---|---|
| PCRISPRi-sgRNA *luxA* | pBBR1-MCS-2 plasmid carrying dcas9 and sgRNA *luxA*, driven by the aTc -inducible TetR promoter, KanR | [43] |
| PCRISPRi-sgRNA *amb0994* | pBBR1-mcs2 plasmid carrying dcas9 and sgRNA *amb0994*, driven by the IPTG-inducible tac promoter, $Kan^R$ | this study |
| pUX19 | Suicide vector, $Kan^R$ | [45] |
| pUCGM | Cloning vector, $Gm^R$ | [45] |
| pUXsuc0994 | pUX19 plasmid carrying HDR DNA, $Kan^R$, $Gm^R$ | this study |
| pBBR1-mcs2 | the broad-host range vector, KanR | [37] |
| pAK 20 | pBBR1-mcs2 plasmid carrying *mamA*-GFP, driven by the IPTG-inducible tac promoter, $Kan^R$ | [46] |
| pAK0994 | pBBR1-mcs2 plasmid carrying *amb0994*-GFP,driven by the IPTG-inducible tac promoter, $Kan^R$ | this study |
| pCRISPR-*amb0994* | pBBR1-mcs2 plasmid carrying dcas9, sgRNA *amb0994* and HDR DNA, driven by the IPTG-inducible tac promoter, $Kan^R$, $Gm^R$ | this study |

**Table S2** Primers used in this study

| Primers | Sequence 5′-3′ | Reference |
|---|---|---|
| sgRNA-0994-F | ggACTAGTGTAATATCGACCATGATTGGGTTTTAGAGCT AGAAATAGCAAGTTAAAATAAGGC | this study |
| sgRNA-0994-R | ccgcTCTAGACAAAAAAAGCACCGACTCGGTGCCACTTT TTCAAGTTGATAACGGACTAGCCTTATTTTAACTTGCTA TTTCTAGCTCTAAAAC | this study |
| sgRNA-Control-F | ggACTAGTGTTTTAGAGCTAGAAATAGCAAGTTAAAAT AAGGC | this study |
| Com *amb0994*-*EcoRI*-GFP-F | ggcGAATTCATGGAAACGACCCTCGGCTCATATG | this study |
| Com *amb0994*-*BamHI*-GFP-R | ggcGGATCCCCGCCCGCCAATTCGGGCGATAAA | this study |
| Cas9-*XhoI*-F1 | ccgCTCGAGATGGATAAGAAATACTCAATAGGCTTAGA TATC | this study |
| Overlap Cas9-R2 | GAAACTTTGTGGAACAATGTGATCGACATC | this study |
| Overlap Cas9-F3 | GATGTCGATCACATTGTTCCACAAAGTTTC | this study |
| Cas9-*XhoI*-R4 | ccgCTCGAGTTAGTCACCTCCTAGCTG | this study |
| 0994-Lift arm-*XbaI*-*XmaI* F1 | gcTCTAGA cccCCCGGGTTCACAAACAGATGCGATAC | this study |

| | | |
|---|---|---|
| 0994-Lift arm-*BamHI* R2 | cgCGGATCCTTTCCAAAGGCTTCAGCG | this study |
| 0994-Right arm-*BamHI* F3 | cgCGGATCC CTGGTCAGGTTGCTGATG | this study |
| 0994-Right arm-*SacII* R4 | tccCCGCGGATGTCCAATAAGCCAAGTCT | this study |
| 1_0994 -F | CAGGTTCCAGGCATTCA | this study |
| 2_0994-delete-R | AATGGTGACAACTACTG | this study |
| 3_Gm-R | AAGCCTGTTCGGTTCGTA | this study |
| Gm-F | CGGCGTTGTGACAATTTAC | this study |
| Check 0995-R | CTTGTCGCCGATCAGAA | this study |
| P1_*mamC*-F | CTGCGATTCCATCATGCGAAAC | this study |
| P1_*mamC*-R | CGATCTGATCGCCATCAACGTC | this study |
| P2_*masA*-F | GCCTGATGGATAGCAACGAAAAAG | this study |
| P2_*masA* -R | AAAGGACGATAAGGCGCAACAG | this study |
| P3_*mamB*-F | GTATCCTGGGCTCCAATCTTGTG | this study |
| P3_*mamB*-R | TGCCGAATACGGCTC AACATAC | this study |
| P4_*mamY*-F | CGGTTCGGAATGGAATGACCATAG | this study |
| P4_*mamY*-R | AGCCTTCGGCAAGTTGAATTCC | this study |
| P5_*mamO′*-F | CCATTCCATCAAGGGACGCTTC | this study |

| | | |
|---|---|---|
| P5_*mamO′*-R | TCTCCTCCACCACGTACAACTG | this study |
| P6_*mamK*-F | CGAACGGAGTGACAAAAATGAGTG | this study |
| P6_*mamK*-R | ACCATGCCACTGTCCTAGACTG | this study |
| Q-*rpoD*-F | ATGGCATCCACCTCAACAAC | [47] |
| Q-*rpoD*-R | CGTAATAGGCGTCGAGGAAG | [47] |
| Q-*amb0994*-F | GCGACAGAATTGGAAGCA | this study |
| Q-*amb0994*-R | CTTGATGGAGGCAGAGAAC | this study |
| Q-*dCas9*-F | GTCGCCGTTATACTGGTT | this study |
| Q-*dCas9*-R | GTCCAGACACTTGTGCTT | this study |

**Movie S1** This movie shows the trajectory of "U turn" after the magnetic field is reversed. The swimming of *M. magneticum* AMB-1 strains control (no sgRNA) was recorded at 33 fps with a 40X objective. The magnetic field was 1 mT in the "U turn" experiment without shielding the geomagnetic field. Red and blue arrows represent the direction of the magnetic field.

**Movie S2** This movie shows the swimming behaviors of KD*amb0994* in the response to a magnetic field.
